# Insulin-like growth factor-2 does not improve behavioral deficits in mouse and rat models of Angelman Syndrome

**DOI:** 10.1101/2021.08.13.456299

**Authors:** Elizabeth L. Berg, Stela P. Petkova, Heather A. Born, Anna Adhikari, Anne E. Anderson, Jill L. Silverman

## Abstract

**Background:** Angelman Syndrome (AS) is a rare neurodevelopmental disorder for which there is currently no cure or effective therapeutic. Since the genetic cause of AS is known to be dysfunctional expression of the maternal allele of ubiquitin protein ligase E3A (*UBE3A*), several genetic animal models of AS have been developed. Both the *Ube3a* maternal deletion mouse and rat models of AS reliably demonstrate behavioral phenotypes of relevance to AS and therefore offer suitable *in vivo* systems in which to test potential therapeutics. One promising candidate treatment is insulin-like growth factor-2 (IGF-2), which has recently been shown to ameliorate behavioral deficits in the mouse model of AS and improve cognitive abilities across model systems.

**Methods:** We used both the *Ube3a* maternal deletion mouse and rat models of AS to evaluate the ability of IGF-2 to improve electrophysiological and behavioral outcomes.

**Results:** Acute systemic administration of IGF-2 had an effect on electrophysiological activity in the brain and on a metric of motor ability, however the effects were not enduring or extensive. Additional metrics of motor behavior, learning, ambulation, and coordination were unaffected and IGF-2 did not improve social communication, seizure threshold, or cognition.

**Limitations:** The generalizability of these results to humans is difficult to predict and it remains possible that dosing schemes (i.e., chronic or subchronic dosing), routes, and/or post-treatment intervals other than that used herein may show more efficacy.

**Conclusions:** Despite a few observed effects of IGF-2, our results taken together indicate that IGF-2 treatment does not profoundly improve behavioral deficits in mice or rat models of AS. These findings shed cautionary light on the potential utility of acute systemic IGF-2 administration in the treatment of AS.

## Background

Angelman Syndrome (AS) is a rare neurodevelopmental disorder caused by the loss of functional ubiquitin protein ligase E3A [1]. Specifically, AS results from deficient expression of the maternal allele, which leaves the entire brain deficient of UBE3A due to neuron-specific imprinting that silences the paternal allele [2–6]. AS is characterized by developmental delay, intellectual disability, impaired communication, gross and fine motor deficits, as well as seizures [7–12]. Since these symptoms are severe and persistent, and there is currently no effective therapeutic or cure for the disorder, those with AS require lifelong supportive care. It is therefore imperative that novel strategies to treat AS are developed.

Several *in vivo* models have been generated to aid in the pursuit of effective treatments, including a conventional germline mouse [13] with a deletion of *Ube3a* in exon 2, a conditional mouse with tamoxifen reactivation [14], a larger deletion mouse [15], and rat model with a full *Ube3a* gene deletion [16]. Various models recapitulate phenotypes of AS and therefore provide useful systems in which to test candidate treatments. Lacking a functional level of UBE3A protein in the brain, models show hypo- locomotion, poor balance, impaired coordination, atypical gait, complex cognitive deficits, alongside communication deficits and aberrant social behavior. Since many of these behavioral deficits are not unique to AS, therapies that are effective for other disorders with shared symptomology, such as autism or other syndromic NDDs, may also be effective in treating AS [17–21].

Insulin-like growth factors (IGFs), a family of proteins with similar structure to insulin, have recently emerged as potential treatments for the social deficits, communication impairments, and repetitive behaviors of genetic syndromes associated with autism spectrum disorder (ASD) [18, 22–30]. IGF-1 is being evaluated as a novel treatment for core symptoms of syndromic autisms in one of the first clinical trials of its kind (NCT01970345) [17-21, 28-33]. IGF-1 is an FDA approved, commercially available compound that crosses the blood-brain barrier and has beneficial effects on synaptic development by promoting neuronal cell survival, synaptic maturation, and synaptic plasticity. Since IGF- 1 has shown efficacy in reversing deficits in mouse and neuronal models of three single gene causes of ASD (namely Rett syndrome [22, 23, 26], Phelan McDermid syndrome [27, 34], and Fragile X syndrome [28]), it may therefore be effective in treating autism spectrum disorders more broadly.

IGF-2, which is important for normal growth and development, tissue repair, and regeneration, has also shown promising effects on ASD-relevant behavioral domains in preclinical studies [35–40]. Injections into the hippocampus have demonstrated that IGF-2 is crucial to the consolidation and enhancement of memories and may be effective in ameliorating memory impairments [30, 41–43]. Since the chemical properties of IGF-2 allow it to exert action within the central nervous system after crossing the blood-brain barrier [44, 45], systemic delivery of IGF-2 represents a highly translational route of treatment. A study in mice by Stern et al. (2014) found that following systemic administration of IGF-2 via subcutaneous injection, adult male C57BL/6J mice showed enhanced novel object recognition, social recognition, contextual fear memory, and working memory [42]. Moreover, in the BTBR mouse model of ASD, Steinmetz et al. (2018) found that IGF-2 treatment normalized behavior in the marble burying task, improved social interaction and social memory deficits, and enhanced novel object recognition along with other types of memory [30].

Despite substantial biological and behavioral differences between the inbred strain BTBR, previously used as an idiopathic ASD model, and the *Ube3a* maternal deletion model of AS, the *Ube3a*^mat-/pat+^ mouse model of AS was recently reported by Cruz et al. (2020) to exhibit behavioral rescue following acute systemic IGF-2 treatment [46]. These encouraging results prompted us to i) investigate if the effects of IGF-2 would be rigorous, reproducible, and inter-laboratory reliable, ii) examine both the mouse and rat model of AS to determine whether IGF-2 could ameliorate or reduce the severity of communication deficits unique to the rat model of AS [16] and evaluate phenotypes observed across species (i.e., motor impairment), and iii) extend the standard, albeit non-translational, rescue of performance in the cerebellar dependent rotarod assay to a rescue of nuanced impairments in gait, which are being utilized as outcome measures in both AS models and AS individuals.

Following a dose range investigation using intra-cranial electroencephalography (EEG) recordings, we employed a battery of behavioral assays to evaluate the effect of systemic IGF-2 on social communication and several motor and learning outcomes in the mouse and rat models of AS. A subcutaneous injection was used to deliver IGF-2 to mice and rats 20 minutes prior to the start of testing. We utilized the standard behavioral protocol of our laboratory and IDDRC behavioral core [16, 47–54] as well as the published protocols of the Alberini laboratory [46] to compare data directly, fairly, and congruently. A comprehensive battery of tests confirmed that IGF-2 did not change basic functions including physical characteristics, general behavioral responses, and sensory reflexes, which indicated safety. Disappointingly, however, our data did not provide strong support for reproducibility or inter-laboratory reliability of IGF-2’s improvement on outcomes since we observed a general lack of effect of IGF-2 in several behavioral domains across two AS rodent models.

## Methods

### Subjects

All animals were housed in a temperature-controlled vivarium and provided food and water *ad libitum*. Animals were maintained on a 12:12 light-dark cycle with the exception of those used for EEG, which were maintained on a 14:10 light-dark cycle. All procedures were approved by the Institutional Animal Care and Use Committee of the University of California, Davis or the Baylor College of Medicine and conducted in accordance with the National Institutes of Health Guide for the Care and Use of Laboratory Animals. Mouse colonies were maintained by breeding *Ube3a* deletion males (B6.129S7- *Ube3a*^tm1Alb^/J; Jackson Laboratory, Bar Harbor, ME; Stock No. 016590) with congenic C57BL/6J (B6J) female mice, and rat colonies were maintained by breeding *Ube3a* deletion males with wildtype Sprague Dawley females (Envigo, Indianapolis, IN). Subject animals were generated by breeding *Ube3a* deletion females with wildtype males, producing maternally inherited *Ube3a* deletion animals (*Ube3a*^mat-/pat*+*^; mat-/pat+; Angelman Syndrome model) and wildtype littermate controls (*Ube3a*^mat+/pat+^; mat+/pat+). Additionally, a mixed-sex cohort of congenic B6J mice was generated from B6J breeder pairs and tested following methods previously described by Cruz et al. (2020) [46] and outlined again in **Supplementary File 1**.

Pups were marked for identification and genotyped as previously described [16, 55]. In order to minimize carry-over effects from repeated testing and handling, at least 24 hours were allowed to elapse between the end of one task and the start of another, and assays were performed in order of least to most stressful. Group sizes for behavioral testing were determined based on previously observed phenotypes and the field recommendation of 10-20 animals for a given task [51]. All behavioral testing included both sexes, was conducted blinded to genotype and treatment group, and was carried out between 08:00 and 18:00 h (ZT1-ZT11) during the light phase. Between subjects, all surfaces of the testing apparatus were cleaned using 70% ethanol and allowed to dry. For assays involving bedding, the bedding was replaced between subjects. At least 1 hour prior to the start of behavioral testing, mice were habituated in their home cages to a dimly lit empty holding room adjacent to the testing area. Two cohorts of mice were tested as follows: Cohort 1 was sampled from 22 litters and, beginning at 8 weeks of age (PND 55), was tested in i) open field, ii) beam walking, iii) DigiGait, iv) novel object recognition, and v) pentylenetetrazol-induced seizures; Cohort 2 was sampled from 15 litters and beginning at 8 weeks of age were tested in i) accelerating rotarod and ii) marble burying. Two cohorts of rats were tested as follows: Cohort 1 was sampled from 6 litters and was tested in i) accelerating rotarod at PND 38 ± 4; Cohort 2 was sampled from 7 litters and was tested in i) pup ultrasonic vocalizations at PND 10 and ii) pro-social USV playback at 9 weeks of age. One mixed-sex cohort of 7 rats was used for recording EEG at 1-2 months of age.

### Systemic treatment with insulin-like growth factor-2 (IGF-2)

IGF-2 (catalog #792-MG, R&D Systems, Inc., Minneapolis, MN) was dissolved in 0.1% bovine serum albumin (BSA) in phosphate-buffered saline (PBS). Prior to testing, a random number generator was used to randomly assign subjects of each genotype to receive either IGF-2 or vehicle (0.1% BSA-PBS). IGF-2 solutions were made fresh prior to every task and, for multi-day tests, injections were carried out only on the first training day. The acute systemic dosing paradigm used herein was based on previous studies showing IGF-2 enhancing cognition [42, 43] and improving behavioral phenotypes of *Ube3a*^mat-/pat+^ mice when administered 20 min prior to testing [46]. Therefore, for all behavioral tests, IGF-2 was delivered 20 min prior to the task. For optimal post-injection data quality while maintaining relevance to the timescale of behavioral tests, IGF-2 was administered 60 min prior to EEG collection. A minimum of two days was allowed to elapse between injections. The 30 µg/kg IGF-2 dose administered to rats was selected based on a dose response analysis of EEG activity following administration of 10, 30, and 60 µg/kg IGF-2 in conjunction with previous data showing efficacy of 30 µg/kg IGF-2 in *Ube3a*^mat-/pat+^ mice [46]. We administered 30 µg/kg IGF-2 to match the dose previously found effective by Cruz et al. [46].

### Electroencephalography (EEG)

To acquire EEG recordings, rats were implanted with two subdural electrodes over the somatosensory cortex and one hippocampal depth electrode as previously described [56]. Rats were anesthetized with isoflurane and positioned within a stereotaxic frame. The cortical recording electrodes were placed at -1.0 mm posterior and ± 3.0 mm lateral relative to bregma, while the hippocampal depth electrode was placed -4.0 mm posterior, +2.8 mm lateral, and -2.8 mm ventral. Metabond (Parkell, Edgewood, NY) and dental cement (Co-Oral-Ite Dental Mfg; Diamond Springs, CA) were used to secure all electrodes, except for the ground electrode which was sutured in the cervical paraspinous region. Electrodes were connected to the commutator via 6-channel pedestal and rats were given minimum 1 week recovery prior to data collection. For pain management, rats were provided with slow release buprenorphine and lidocaine/bupivacaine on the day of surgery, as well as Rimadyl tablets on the day prior to, the day of, and the day after surgery. Video synchronized EEG data was acquired using the Nicolet system (Natus, Pleasanton, CA) and Labchart V8 software (AD Instruments, Colorado Springs, CO) and then inspected and analyzed by a trained experimenter blinded to genotype and treatment group. Pre-injection baseline data (60 min in duration) were recorded from rats 24 hrs prior to administration of vehicle and post-injection data (60 min in duration) were collected 60 min following injection. Data were analyzed using repeated measures ANOVA with group as the between-group factor and frequency as the within-group factor or using two-way ANOVA with genotype and IGF-2 treatment as between-group factors.

### Behavioral Assays

#### Accelerating rotarod

To test motor coordination, balance, and motor learning, subjects were placed on an Ugo-Basile accelerating rotarod (Stoelting Co., Wood Dale, IL) as described previously in mice and rats [16, 47, 57]. Animals were placed on the cylinder while it rotated at 5 revolutions per minute, and then it slowly accelerated to 40 revolutions per min over the course of the 5 min trial. On three consecutive days, subjects were given three trials per day with a 45-60 min inter-trial rest period. The latency for each subject to fall off the cylinder was recorded with the maximum achievable latency being 300 seconds. Data were analyzed using three-way ANOVA with genotype and treatment as the between-group factors and day as the within-group factor.

#### Isolation-induced pup ultrasonic vocalizations (USV)

On PND 10, neonatal rats were assessed by collecting 40 kHz vocalizations made when isolated from dam and littermates following a previously described protocol [16, 47, 57, 58]. Rat pups were selected from the nest at random and placed in a small plastic container with clean bedding. The container was placed inside a sound attenuating chamber for three min while calls were recorded with an ultrasonic microphone and Avisoft-RECORDER software (Avisoft Bioacoustics, Glienicke, Germany). Using spectrograms generated with Avisoft-SASLab Pro software, calls were manually counted by a trained investigator blinded to genotype and treatment group. Data were analyzed using two-way ANOVA with genotype and IGF-2 treatment as between-group factors.

#### Pro-social USV playback

To evaluate social behavior, the behavioral response to hearing playback of natural conspecific 50-kHz USV social contact calls was quantified following an established protocol [16, 47, 59]. Prior to the test, all subjects were handled in a standardized manner for 5 min on three consecutive days. Subjects were individually placed on an eight-arm radial maze (arms: 40 cm l x 10 cm w) elevated 48 cm above the floor, surrounded by a black curtain, and illuminated to ∼8 lux with indirect white light. An active ultrasonic speaker (ScanSpeak, Avisoft Bioacoustics) was placed 20 cm away from the end of one arm while a second inactive speaker was placed symmetrically at the opposite arm to serve as a visual control. After a 15-min habituation period, an Ultra SoundGate 116 Player (Avisoft Bioacoustics) was used to present one of two 1-min acoustic stimuli: (1) pro-social 50-kHz USV or (2) a time- and amplitude-matched white noise control stimulus. Following a 10-min inter-stimulus interval, the second stimulus was presented, and the test session ended after an additional 10-min post-stimulus period. The order of the stimuli was counterbalanced in order to account for possible sequence effects. An overhead camera and EthoVision XT videotracking software (Noldus Information Technology, Wageningen, Netherlands) were used to measure stimulus-induced changes in locomotion and location on the maze. Intact behavioral inhibition was defined as moving significantly less during the minute of white noise compared to the minute prior by paired *t-*test. Intact social approach was defined as spending significantly more time on the arms proximal to the active speaker compared to the distal arms during the minute of pro-social 50-kHz USV playback and subsequent two min by paired *t*-test. As a control metric for motor behavior, distance traveled during this timeframe (i.e., the minute of USV playback and subsequent two min) was also analyzed using two-way ANOVA with genotype and IGF-2 treatment as between-group factors.

#### Open field locomotion

General exploratory locomotion was assayed as previously described [55, 60, 61]. Subjects were individually placed within a novel open field (40 cm l x 40 cm w x 30.5 cm h), which was dimly illuminated to ∼30 lux, and allowing them to explore for 30 min. Photocell beam breaks were detected automatically by the VersaMax Animal Activity Monitoring System (AccuScan Instruments, Columbus, OH) to measure horizontal activity, vertical activity, and center time. Data were analyzed using two-way ANOVA with genotype and IGF-2 treatment as between-group factors.

#### Beam walking

A beam walking motor task was carried out by individually placing subjects at one end of a 59 cm long beam as described previously [60]. The beam was elevated 68 cm above a cushion and the time taken to cross the beam was measured. A darkened goal box (12 cm d cylinder) was placed on the far end of the beam in order to provide motivation to walk across. On the first day, three training trials on a large diameter (35 mm) beam were conducted to allow animals to become accustomed to the task. Animals that had scores of 60 seconds on all three trials were excluded from analysis. On the following day, subjects were placed back on the large diameter beam and then on a beam of intermediate width (18 mm d) before being placed onto the test beam, which was the narrowest and therefore most challenging (13 mm d). Two trials per beam were carried out with an inter-trial rest interval of at least 30 minutes and trial duration was capped at a maximum of 60 seconds. The two-trial average latency to traverse the test beam was recorded and data were analyzed via two-way ANOVA with genotype and IGF-2 treatment as between-group factors.

#### Marble burying

To evaluate marble burying, twenty black glass marbles (15 mm d) were arranged in a 4 x 5 grid on top of 4 cm of clean bedding within a standard mouse cage (27 cm l x 16.5 cm w x 12.5 cm h) following a protocol similar to those described previously [49, 62]. Subjects were individually placed in the center of the cage and allowed to explore for 20 min. The testing room was dimly illuminated to ∼15 lux. The number of marbles buried (defined as at least 50% covered by the bedding) at the end of the test session was recorded. Data were analyzed using two-way ANOVA with genotype and IGF-2 treatment as between-group factors.

#### Pentylenetetrazol-induced seizures

Susceptibility to primary generalized seizures was behaviorally assessed by systemically administering 80 mg/kg pentelenetetrazol (PTZ; a GABA_A_ receptor antagonist) via intraperitoneal injection and observing the timing and progression of the subsequent convulsions following a protocol described previously [55, 63, 64]. Immediately following injection of PTZ, animals were individually placed in a clean empty standard mouse cage (27 cm l x 16.5 cm w x 12.5 cm h) and watched carefully by a trained observer blinded to genotype and treatment condition. The latency to generalized clonus was recorded and analyzed using two-way ANOVA with genotype and IGF-2 treatment as between-group factors.

#### Novel object recognition (NOR)

Learning and memory were tested by individually presenting subjects with two identical objects and later testing their ability to recognize the familiar object over a novel one using an established protocol previously described [49, 53, 60, 65]. The NOR assay was carried out within an opaque matte white arena (41 cm l x 41 cm w x 30 cm h) in a 30-lux room and consisted of five phases: a 30-min habituation to the arena on the day prior to the test, a 10-min habituation to the arena on the test day, a 10-min object familiarization session, a 60-min isolation period, and a 5-min object recognition test. Following the 10-min habituation on the day of the test, each animal was removed from the arena and placed in an individual clean holding cage while two clean identical objects were placed inside the arena. Each subject was then returned to its arena and allowed to explore and familiarize with the objects for 10 min. Subjects were then returned to their holding cages and placed in a nearby low light holding area outside of the testing room. The arenas were cleaned, let dry, and one clean familiar object and one clean novel object were placed inside the arena where the two identical objects had previously been located. After a 60 min interval, subjects were returned to their arenas and allowed to explore the objects for 5 min. Time spent investigating each object was measured using EthoVision XT videotracking software (Noldus Information Technology) and validated by manual scoring by a trained observer blinded to genotype and treatment group. Object investigation was defined as time spent sniffing the object when the nose was within 2 cm of the object and oriented toward the object. Animals who did not spend at least 5 sec sniffing the objects during the familiarization phase were removed from analysis and recognition memory was defined as spending significantly more time investigating the novel object compared to the familiar object by paired *t*-test within group. Object preference was calculated as time spent sniffing the novel object compared to total time sniffing both objects. Fifty percent represents equal time investigating the novel and familiar object (a lack of preference) whereas >50% demonstrates intact recognition memory.

#### DigiGait

Gait metrics were collected using the DigiGait automated treadmill system and analysis software (Mouse Specifics, Inc., Framingham, MA). Subjects were placed individually into the enclosed treadmill chamber and allowed to acclimate before the belt was turned on and the speed was slowly increased from 5 cm/sec to a constant speed of 20 cm/sec. For each subject, 3-6 sec of clearly visible consecutive strides at the belt speed of 20 cm/sec was recorded. Gait analysis was conducted using the DigiGait software package and was carried out by an experimenter blinded to genotype and treatment condition. Right and left fore- and hindlimbs were averaged together. Data were analyzed per limb set using two-way ANOVA with genotype and IGF-2 treatment as between-group factors.

#### Statistical Analysis

All statistical analyses were carried out using Prism 9 software (GraphPad Software, San Diego, CA). All significance levels were set at *p* < 0.05 and all *t*-tests were two-tailed. Outliers were identified and excluded using Grubb’s test and D’Agostino & Pearson tests were used to check assumptions of normality. Two-way ANOVAs were used to analyze the effects of both genotype and IGF-2 treatment and two-way repeated measures ANOVAs were used for comparisons across time points. Three-way ANOVAs were used to analyze the effects of genotype, treatment, as well as time. Paired *t*- tests were used for comparisons within a single group. Subsequent to ANOVAs, *post hoc* testing controlling for multiple comparisons was carried out using Sidak’s or Tukey’s multiple comparisons test. Since the overall goal of the study was to evaluate the potential for IGF-2 to ameliorate behavioral deficits, emphasis was placed on i) the comparison between wildtype *Ube3a*^mat+/pat+^ vehicle and *Ube3a*^mat-/pat+^ vehicle to confirm the genotype deficit and ii) the comparison between *Ube3a*^mat-/pat+^ vehicle and *Ube3a*^mat-/pat+^ IGF-2 to identify any effect of IGF-2 treatment on the deficit. Data are presented as mean ± standard error of the mean (S.E.M.) unless otherwise noted and detailed statistics are described in **Supplementary File 2**. No significant sex differences were detected so data from both sexes were pooled together.

## Results

### IGF-2 reduced cortical and hippocampal delta power in in *Ube3a*^mat-/pat+^ rats

Since *Ube3a*^mat-/pat+^ rats display the elevation in EEG delta power that is characteristic of AS [56], we sought to examine whether this core phenotype could be normalized by IGF-2. Prior to treatment, we used cortical and hippocampal electrodes to conduct spectral power analyses in *Ube3a*^mat-/pat+^ rats, which revealed elevations in the delta range (1-4 Hz), although when analyzed across the entire frequency range, the effect of genotype was not statistically significant (**Fig. 1A**; *F*_Genotype_, *p*>0.05; *F*_Frequency_, *p*<0.0001; *F*_G×F_, *p*>0.05; **Fig. 1B**; *F*_Genotype_, *p*>0.05; *F*_Frequency_, *p*<0.0001; *F*_G×F_, *p*>0.05). In wildtype rats, treatment with IGF-2 did not influence cortical power (**Fig. 1C**; *F*_Genotype_, *p*>0.05; *F*_Frequency_, *p*<0.0001; *F*_G×F_, *p*>0.05) or hippocampal power (**Fig. 1D**; *F*_Genotype_, *p*>0.05; *F*_Frequency_, *p*<0.0001; *F*_G×F_, *p*>0.05). In *Ube3a*^mat-/pat+^ rats, however, treatment with IGF-2 reduced cortical power at 1, 2, 3, and 4 Hz (**Fig. 1E**; *F*_Genotype_, *p*>0.05; *F*_Frequency_, *p*<0.0001; *F*_G×F_, *p*<0.0001). At 1 and 2 Hz, all doses of IGF-2 reduced cortical delta power in *Ube3a*^mat-/pat+^ compared to vehicle (10 µg/kg IGF-2, *p*<0.0001; 30 µg/kg IGF-2, *p*<0.0001; 60 µg/kg IGF-2, *p*<0.0001). At 3 Hz, cortical delta power was reduced by 10 µg/kg IGF-2 (*p*<0.0001), 30 µg/kg IGF-2 (*p*<0.0001), and 60 µg/kg IGF-2 (*p*=0.027) and was reduced at 4 Hz by 10 µg/kg IGF-2 (*p*=0.002) and 30 µg/kg IGF-2 (*p*=0.021) but not by 60 µg/kg IGF-2 (*p*>0.05). Despite trending reductions, hippocampal delta power was not affected by IGF-2 treatment in *Ube3a*^mat-/pat+^ rats (**Fig. 1F**;; *F*_Genotype_, *p*>0.05; *F*_Frequency_, *p*<0.0001; *F*_G×F_, *p*>0.05).

**Fig. 1.**
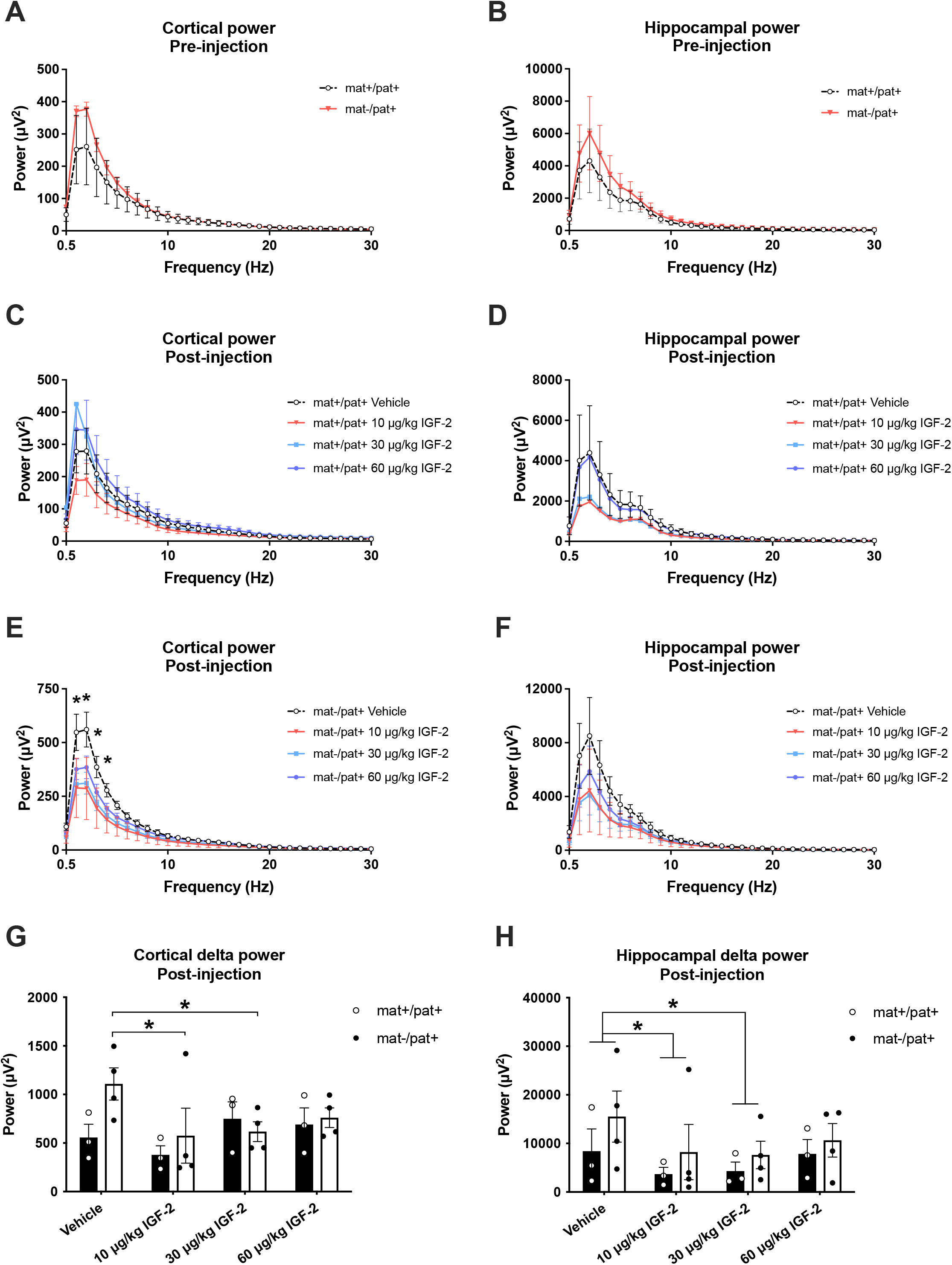
IGF-2 reduced cortical and hippocampal delta power in *Ube3a*^mat-/pat+^ rats. **(A)** Baseline cortical and **(B)** hippocampal power pre-injection trended higher in *Ube3a*^mat-/pat+^ (mat-/pat+) rats compared to wildtype (*Ube3a*^mat+/pat+^; mat+/pat+) rats. **(C)** Following injection of IGF-2, cortical and **(D)** hippocampal power was unaltered in wildtype rats. **(E)** In mat-/pat+ rats, treatment with 10 or 30 µg/kg IGF-2 led to reduced cortical power at 1-4 Hz, while treatment with 60 µg/kg IGF-2 reduced cortical power at 1-3 Hz. **(F)** Hippocampal power was unchanged by IGF-2 in mat-/pat+ rats. **(G)** Cortical power at 1 and 2 Hz (“delta power”) was lower in mat-/pat+ rats following treatment with 10 or 30 µg/kg IGF-2 compared to vehicle. **(H)** Across both genotypes, hippocampal delta power (at 1 and 2 Hz) was also reduced by 10 and 30 µg/kg IGF-2. Data are expressed as mean ± S.E.M. *n*=3-4 rats/genotype. E: \**p*<0.05 vs. mat-/pat+ Vehicle, Sidak’s multiple comparisons following repeated measures ANOVA. G, H: \**p*<0.05, Tukey’s multiple comparisons following repeated measures ANOVA.

To more closely examine dose differences on the EEG phenotype of *Ube3a*^mat-/pat+^ rats, we analyzed each dose’s effect on summed power at 1 and 2 Hz (“delta power”). These two frequencies were of main interest due to peak signal strength across our spectral analyses as well as previous work that identified 1-2 Hz as showing the most persistent difference between *Ube3a*^mat-/pat+^ and wildtype rats [56]. We found no effect of IGF-2 on delta power in wildtype but cortical delta power in *Ube3a*^mat-/pat+^ rats was reduced following treatment with 10 or 30 µg/kg IGF-2 (**Fig. 1G**; *F*_Genotype_, *p*>0.05; *F*_Treatment_, *p*=0.031; *F*_G×F_, *p*=0.039; IGF-2 vs. vehicle, 10 µg/kg, *p*=0.008; 30 µg/kg, *p*=0.015; 60 µg/kg, *p*>0.05). Hippocampal delta power did not differ by genotype but was reduced by treatment with 10 or 30 µg/kg IGF-2 (**Fig. 1H**; *F*_Genotype_, *p*>0.05; *F*_Treatment_, *p*=0.027; *F*_G×F_, *p*>0.05; IGF-2 vs. vehicle, 10 µg/kg, *p*=0.045; 30 µg/kg, *p*=0.045; 60 µg/kg, *p*>0.05). Overall, both 10 and 30 µg/kg IGF-2 showed promising effects to reduce the elevated delta power of *Ube3a*^mat-/pat+^ rats in both the cortex and hippocampus. In selecting a dose to investigate in subsequent behavioral testing of rats, we also considered the demonstrated efficacy of 30 µg/kg IGF-2 in previous studies of IGF-2 [46] and therefore opted to use this dose in rats moving forward.

### IGF-2 did not improve motor learning or social communication in *Ube3a*^mat-/pat+^ rats

In order to assess whether IGF-2 could ameliorate the robust motor learning deficit of *Ube3a*^mat-/pat+^ rats, we tested *Ube3a*^mat-/pat+^ and wildtype littermate controls (*Ube3a*^mat+/pat+^) with IGF-2 or vehicle treatment on an accelerating rotarod (**Fig. 2A**). The motor learning deficit of *Ube3a*^mat-/pat+^ rats was apparent across the three day task, although it was unaffected by treatment with IGF-2 (*F*_Genotype_, *p*=0.038; *F*_Treatment_, *p*>0.05; *F*_Time_, *p*<0.0001; *F*_G×Tr_, *p*>0.05; *F*_G×Ti_, *p*<0.0001; *F*_Tr×Ti_, *p*>0.05; *F*_G×Ti×Tr_, *p*>0.05). While the wildtype vehicle and wildtype IGF-2 groups significantly improved their performance from session 1 to 3 (by 142% and 118%, respectively), both the *Ube3a*^mat-/pat+^ vehicle and *Ube3a*^mat-/pat+^ IGF-2 groups failed to improve over the course of the test (*Ube3a*^mat+/pat+^ vehicle, *p*=0.003; *Ube3a*^mat+/pat+^ IGF-2, *p*=0.002; *Ube3a*^mat-/pat+^ vehicle, *p*>0.05; *Ube3a*^mat-/pat+^ IGF-2, *p*>0.05), in contrast to the recent report by Cruz et al. [46].

**Fig. 2.**
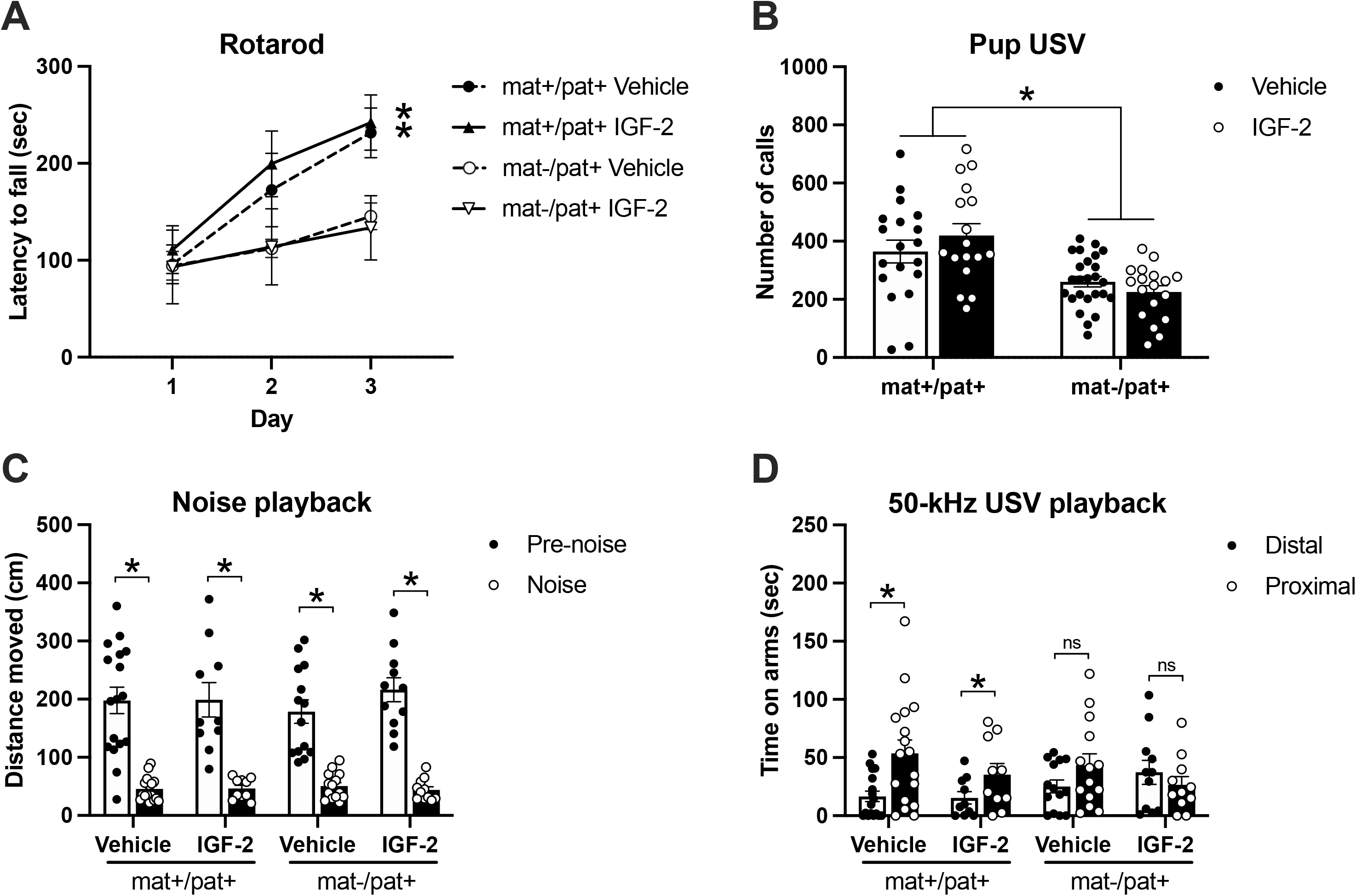
IGF-2 did not rescue or improve motor learning or social communication in *Ube3a*^mat-/pat+^ rats. **(A)** Latency to fall off an accelerating rotarod significantly improved from session 1 to 3 for both wildtype groups (*Ube3a*^mat+/pat+^; mat+/pat+), but not for either *Ube3a*^mat-/pat+^ group (mat-/pat+). **(B)** At PND 10, mat-/pat+ pups emitted fewer isolation-induced ultrasonic vocalizations (USV) than wildtype littermates, but IGF-2 had no effect on vocalization rates. **(C)** All groups showed behavioral inhibition (i.e., reduced locomotion) during playback of white noise compared to baseline. **(D)** During playback of pro-social 50-kHz USV, only the wildtype groups, and not mat-/pat+ rats, spent significantly more time on the arms proximal to the speaker compared to the distal arms (i.e., social approach). Data are expressed as mean ± S.E.M. *n*=6-25 rats/group. A: \**p*<0.05, Day 1 vs. 3, Tukey’s multiple comparisons following three-way ANOVA. B: \**p*<0.05, main effect of genotype, two-way ANOVA. C, D: \**p*<0.05, paired *t*-test. ns, not significantly different, *p*>0.05.

We also evaluated the effect of IGF-2 on social communication outcomes, both at an early postnatal timepoint and during adulthood. *Ube3a*^mat-/pat+^ rats emitted 37% fewer isolation-induced pup USV at PND 10 compared to wildtype, reproducing our earlier publication [16], but IGF-2 had no effect on the calling rate (**Fig. 2B**; *F*_Genotype_, *p*<0.0001; *F*_Treatment_, *p*>0.05; *F*_G×T_, *p*>0.05). Then in adulthood, we used a USV playback paradigm to present subjects with pro-social 50-kHz USV and a time- and amplitude-matched white noise acoustic control (**Fig. 2C**). All groups, regardless of genotype or treatment, exhibited the expected behavioral inhibition in response to the noise control wherein they moved less during playback of the noise compared to pre-noise baseline exploration, indicating intact hearing abilities (*Ube3a*^mat+/pat+^ vehicle, *p*<0.001; *Ube3a*^mat+/pat+^ IGF-2, *p*<0.001; *Ube3a*^mat-/pat+^ vehicle, *p*<0.0001; *Ube3a*^mat-/pat+^ IGF-2, *p*<0.0001). This is further supported by the observation of equivalent levels of locomotion in all groups following initiation of 50-kHz USV playback (data not shown; two-way ANOVA: *F*_Genotype_, *p*>0.05; *F*_Treatment_, *p*>0.05; *F*_G×T_, *p*>0.05). In response to the pro-social 50-kHz USV, only the wildtype vehicle and wildtype IGF-2 groups showed the typical social approach response by spending more time on the arms proximal to the speaker compared to the distal arms (**Fig. 2D**; *Ube3a*^mat+/pat+^ vehicle, *p*=0.004; *Ube3a*^mat+/pat+^ IGF-2, *p*=0.045). Both the *Ube3a*^mat-/pat+^ vehicle and *Ube3a*^mat-/pat+^ IGF-2 groups failed to show a preference for the proximal arms in response to the USV (*Ube3a*^mat-/pat+^ vehicle, *p*>0.05; *Ube3a*^mat-/pat+^ IGF-2, *p*>0.05), reproducing our earlier publication [16]. Given no differences in the response to a non-social stimulus and the absence of a motor impairment, the reduced social approach response in both groups of *Ube3a*^mat-/pat+^ rats reveals a social communication deficit that is not ameliorated by treatment with IGF-2.

### IGF-2 did not markedly improve motor deficits, seizure threshold, or object recognition in *Ube3a*^mat-/pat+^ mice

Next, we examined the ability of IGF-2 to improve the known behavioral deficits of *Ube3a*^mat-/pat+^ mice. While *Ube3a*^mat-/pat+^ mice showed strong motoric deficits, performance was not affected by treatment with IGF-2. First, exploration of a novel open arena was used to assess overall locomotive activity. Horizontal activity, which was 44% lower in *Ube3a*^mat-/pat+^ mice than wildtype littermates, was not affected by IGF-2 (**Fig. 3A**; *F*_Genotype_, *p*<0.0001; *F*_Treatment_, *p*>0.05; *F*_G×T_, *p*>0.05). A similar pattern was observed for vertical activity wherein *Ube3a*^mat-/pat+^ mice showed 60% less rearing and vertical movement compared to wildtype, but this was unaffected by IGF-2 (**Fig. 3B**; *F*_Genotype_, *p*<0.0001; *F*_Treatment_, *p*>0.05; *F*_G×T_, *p*>0.05). There was no genotype difference or effect of IGF-2 on time spent in the center of the open field (**Fig. 3C**; *F*_Genotype_, *p*>0.05; *F*_Treatment_, *p*>0.05; *F*_G×T_, *p*>0.05). These data were similar to the recent Cruz et al. report [46].

**Fig. 3.**
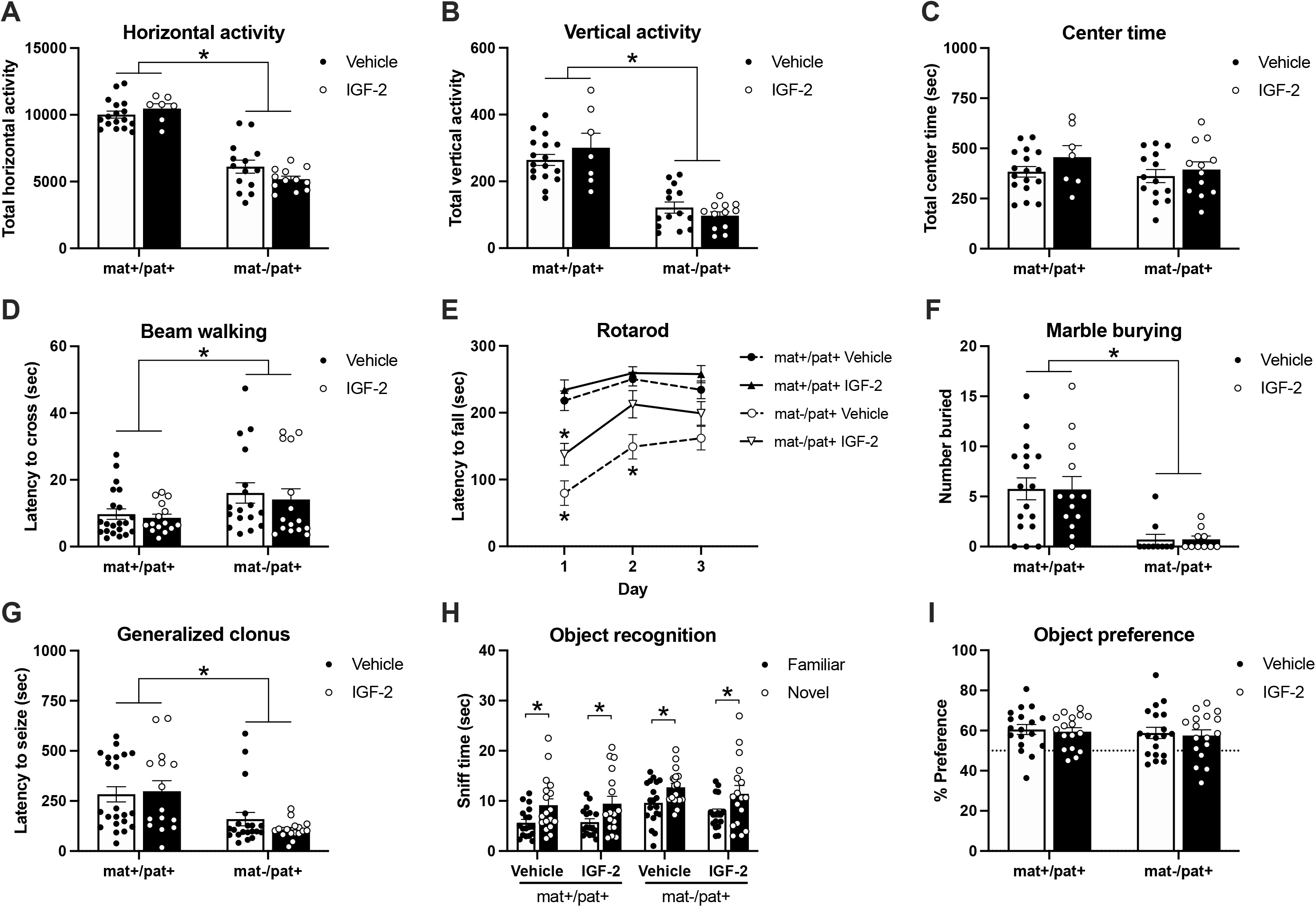
IGF-2 did not markedly improve motor deficits, seizure threshold, or object recognition in *Ube3a*^mat-/pat+^ mice. **(A)** Horizontal and **(B)** vertical activity in an open field assay were reduced in *Ube3a*^mat-/pat+^ mice (mat-/pat+) compared to wildtype littermates (*Ube3a*^mat+/pat+^; mat+/pat+), but unaffected by IGF-2. **(C)** Center time did not differ among groups. **(D)** Latency to cross a thin beam was elevated in mat-/pat+ mice, but unchanged by IGF-2. **(E)** Accelerating rotarod performance was moderately improved by IGF-2 treatment in mat-/pat+ mice, however only on the first day of testing. **(F)** Regardless of IGF-2 treatment, mat-/pat+ mice demonstrated a marble burying deficit and **(G)** mat-/pat+ mice were quicker to exhibit generalized clonus following pentylenetetrazol administration, which was unaffected by IGF-2. **(H)** All groups demonstrated intact novel object recognition as measured by more time spent investigating the novel object compared to the familiar object and by **(I)** novel object percent preference. Data are expressed as mean ± S.E.M. *n*=10-22 mice/group. A-D, F, G: \**p*<0.05, main effect of genotype, two-way ANOVA. E: \**p*<0.05 vs. mat+/pat+ vehicle, Tukey’s multiple comparisons test following three-way ANOVA. H: \**p*<0.05, paired *t*-test.

We also assessed balance and motor coordination using a beam walking task but found that IGF-2 did not have an enhancing effect in wildtypes nor ameliorated motor coordination deficits observed in the *Ube3a*^mat-/pat+^ group. *Ube3a*^mat-/pat+^ mice took longer to cross compared to wildtype littermates regardless of treatment with IGF-2 (**Fig. 3D**; *F*_Genotype_, *p*=0.016; *F*_Treatment_, *p*>0.05; *F*_G×T_, *p*>0.05). However, in the accelerating rotarod task of motor coordination, we were able to detect a moderate effect of IGF-2 in *Ube3a*^mat-/pat+^ mice (**Fig. 3E**; *F*_Genotype_, *p*<0.0001; *F*_Treatment_, *p*=0.006; *F*_Time_, *p*<0.0001; *F*_G×Tr_, *p*>0.05; *F*_G×Ti_, *p*=0.007; *F*_Tr×Ti_, *p*>0.05; *F*_G×Ti×Tr_, *p*>0.05). While *Ube3a*^mat-/pat+^ mice had poorer performance than wildtypes, falling off earlier on test days 1 and 2 (*Ube3a*^mat+/pat+^ vehicle vs. *Ube3a*^mat-/pat+^ vehicle, day 1, *p*<0.001; day 2, *p*=0.008; day 3, *p*>0.05), *Ube3a*^mat-/pat+^ mice treated with IGF-2 only showed a deficit on the first day of testing (*Ube3a*^mat+/pat+^ vehicle vs. *Ube3a*^mat-/pat+^ IGF-2, day 1, *p*=0.047; day 2, *p*>0.05; day 3, *p*>0.05). The effect, however, was only moderate in that the *Ube3a*^mat-/pat+^ IGF-2 group was not significantly better than the *Ube3a*^mat-/pat+^ vehicle group on any day (*Ube3a*^mat-/pat+^ vehicle vs. *Ube3a*^mat-/pat+^ IGF-2, day 1, *p*>0.05; day 2, *p*>0.05; day 3, *p*>0.05).

In the marble burying assay, *Ube3a*^mat-/pat+^ mice covered 88% fewer marbles compared to wildtype littermates but there was no effect of IGF-2 treatment in either group ((**Fig. 3F**; *F*_Genotype_, *p*<0.0001; *F*_Treatment_, *p*>0.05; *F*_G×T_, *p*>0.05). As described in previous reports, our laboratory interprets the lack of marble burying as a function of the low motor activity of AS mice, as opposed to traditional interpretations of anxiety-like or repetitive behavior used by other AS laboratories [66]. In a fully capable, typically active mouse, marble burying may hold more meaning, however, after more than five years of focused study on these mice, we cannot delineate the motor impairments related to marble burying. We also investigated IGF-2’s influence on seizure threshold in *Ube3a*^mat-/pat+^ mice using the chemo-convulsant pentelenetetrazol. While *Ube3a*^mat-/pat+^ mice exhibited a reduced latency to generalized clonus seizure, latency to seize was unaffected by IGF-2 treatment (**Fig. 3G**; *F*_Genotype_, *p*>0.0001; *F*_Treatment_, *p*>0.05; *F*_G×T_, *p*>0.05). *Ube3a*^mat-/pat+^ mice were 53% quicker to seize than wildtype.

To test the cognition enhancing capabilities of IGF-2 treatment, we evaluated novel object recognition with a standard protocol and found that all groups, regardless of genotype or treatment, demonstrated intact novel object recognition (**Fig. 3H**). Within each group, more time was spent more time investigating the novel object compared to the familiar one (*Ube3a*^mat+/pat+^ vehicle, *p*<0.001; *Ube3a*^mat+/pat+^ IGF-2, *p*=0.001; *Ube3a*^mat-/pat+^ vehicle, *p*=0.006; *Ube3a*^mat-/pat+^ IGF-2, *p*=0.006). In addition to the dichotomous yes/no analysis of object recognition, we also explored whether IGF-2 influenced the continuous metric of object preference. There were no differences, however, in percent preference for the novel object across genotypes or treatment (**Fig. 3I**; *F*_Genotype_, *p*>0.05; *F*_Treatment_, *p*>0.05; *F*_G×T_, *p*>0.05). To facilitate more direct comparisons with the results of Cruz et al. (2020), we also utilized their novel object recognition protocol within our own laboratory. We found, however, that IGF-2 failed to elicit recognition memory in congenic C57BL/6J mice, the background strain of the *Ube3a*^mat-/pat+^ mouse model (**Fig. S1**). Concomitantly, using the experimental paradigm of Cruz et al. [46], we observed that IGF-2 treatment did not affect the cognitive performance of C57BL/6J mice in the delayed contextual fear conditioning task (Fig. S1).

As an innovative and unique investigation of nuanced motor phenotypes, we probed for any effect of IGF-2 on several metrics of gait using the automated DigiGait system. While walking on a treadmill, *Ube3a*^mat-/pat+^ mice took wider, longer, and fewer steps compared to wildtype littermates. The elevated forelimb and hindlimb stance widths exhibited by *Ube3a*^mat-/pat+^ mice were not affected by IGF-2 treatment (**Fig. 4A**; fore: *F*_Genotype_, *p*<0.0001; *F*_Treatment_, *p*=0.046; *F*_G×T_, *p*>0.05; *Ube3a*^mat+/pat+^ vehicle vs. *Ube3a*^mat-/pat+^ vehicle, *p*=0.005; *Ube3a*^mat-/pat+^ vehicle vs. *Ube3a*^mat-/pat+^ IGF-2, *p*>0.05; hind: *F*_Genotype_, *p*<0.001; *F*_Treatment_, *p*>0.05; *F*_G×T_, *p*>0.05). Additionally, the longer forelimb and hindlimb stride lengths were further increased by IGF-2 (**Fig. 4B**; fore: *F*_Genotype_, *p*<0.0001; *F*_Treatment_, *p*=0.031; *F*_G×T_, *p*=0.038; *Ube3a*^mat+/pat+^ vehicle vs. *Ube3a*^mat-/pat+^ vehicle*, p*<0.001; *Ube3a*^mat-/pat+^ vehicle vs. *Ube3a*^mat-/pat+^ IGF-2, *p*=0.021; hind: *F*_Genotype_, *p*<0.0001; *F*_Treatment_, *p*>0.05; *F*_G×T_, *p*=0.023; *Ube3a*^mat+/pat+^ vehicle vs. *Ube3a*^mat-/pat+^ vehicle*, p*<0.0001; *Ube3a*^mat-/pat+^ vehicle vs. *Ube3a*^mat-/pat+^ IGF-2, *p*=0.031). IGF-2 also led to further reduction of forelimb stride frequency and did not have an effect on the reduced hindlimb stride frequency displayed by *Ube3a*^mat-/pat+^ mice (**Fig. 4C**; fore: *F*_Genotype_, *p*<0.0001; *F*_Treatment_, *p*>0.05; *F*_G×T_, *p*>0.05; *Ube3a*^mat+/pat+^ vehicle vs. *Ube3a*^mat-/pat+^ vehicle*, p*<0.001; *Ube3a*^mat-/pat+^ vehicle vs. *Ube3a*^mat-/pat+^ IGF-2, *p*=0.021; hind: *F*_Genotype_, *p*<0.0001; *F*_Treatment_, *p*>0.05; *F*_G×T_, *p*>0.05). Interestingly, IGF-2 had varying effects on the time taken to propel each step: the elevated propulsion time required by *Ube3a*^mat-/pat+^ mice, indicative of limb weakness, was unaffected by IGF-2 in the forelimbs while further elevated by IGF-2 in the hindlimbs, whose function is largely force generation and propulsion (**Fig. 4D**; fore: *F*_Genotype_, *p*<0.0001; *F*_Treatment_, *p*>0.05; *F*_G×T_, *p*>0.05; hind: *F*_Genotype_, *p*<0.0001; *F*_Treatment_, *p*>0.05; *F*_G×T_, *p*=0.014; *Ube3a*^mat+/pat+^ vehicle vs. *Ube3a*^mat-/pat+^ vehicle*, p*=0.003; *Ube3a*^mat-/pat+^ vehicle vs. *Ube3a*^mat-/pat+^ IGF-2, *p*=0.032). In alignment with taking longer steps, *Ube3a*^mat-/pat+^ mice held their fore and hindlimbs in a swing state off the ground longer than wildtypes, although neither metric was changed by IGF-2 treatment (**Fig. 4E**; fore: *F*_Genotype_, *p*<0.0001; *F*_Treatment_, *p*>0.05; *F*_G×T_, *p*>0.05; hind: *F*_Genotype_, *p*<0.0001; *F*_Treatment_, *p*>0.05; *F*_G×T_, *p*>0.05). Finally, despite increased propulsion and swing times, *Ube3a*^mat-/pat+^ mice spent a normal amount of time braking, which was unchanged by IGF-2 treatment (**Fig. 4F**; fore: *F*_Genotype_, *p*>0.05; *F*_Treatment_, *p*>0.05; *F*_G×T_, *p*>0.05; hind: *F*_Genotype_, *p*>0.05; *F*_Treatment_, *p*>0.05; *F*_G×T_, *p*>0.05).

**Fig. 4.**
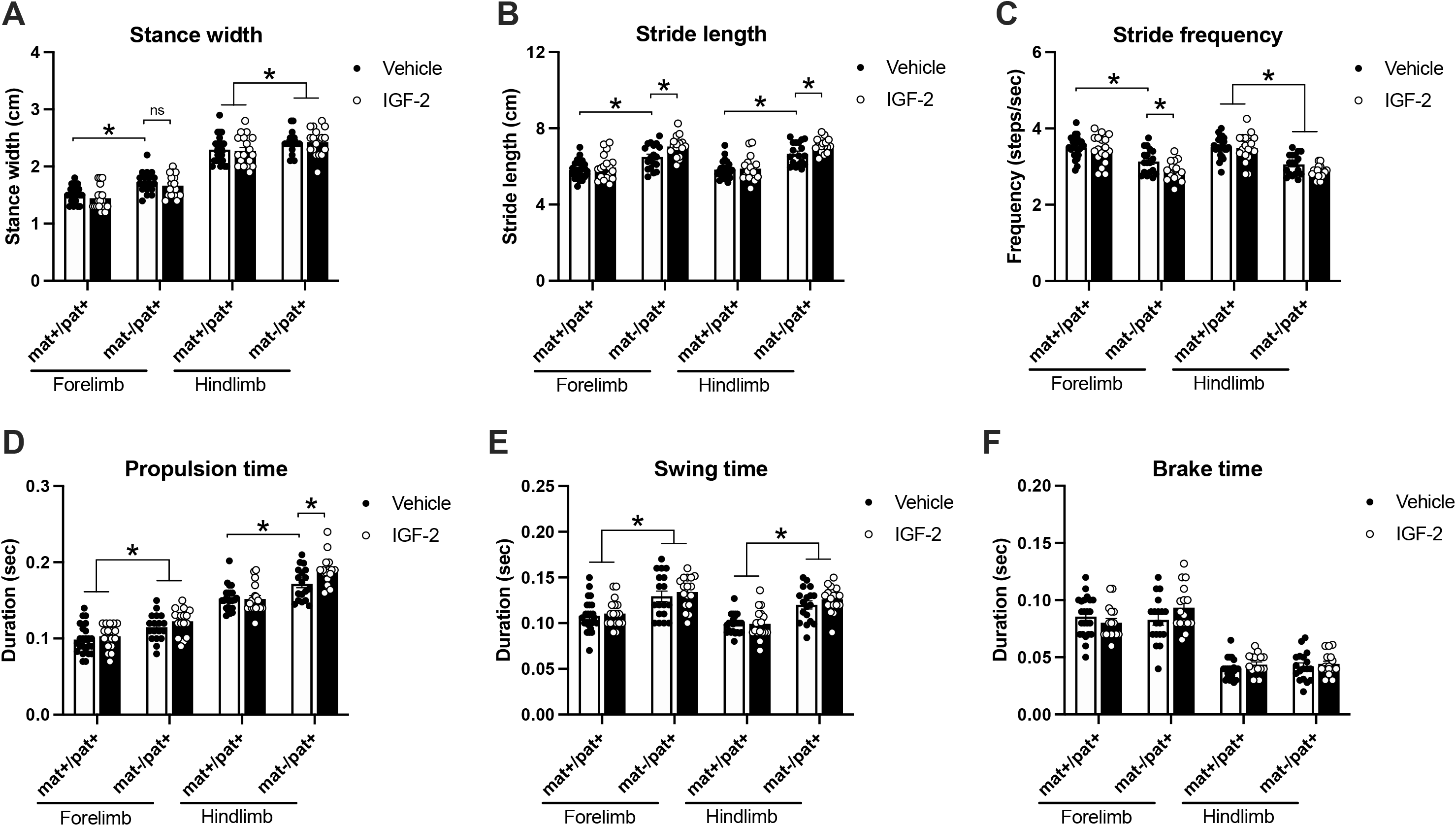
IGF-2 did not rescue or improve gait deficits in *Ube3a*^mat-/pat+^ mice. **(A)** Compared to wildtype littermates (*Ube3a*^mat+/pat+^; mat+/pat+), *Ube3a*^mat-/pat+^ (mat-/pat+) mice exhibited wider stances while treadmill walking, which were unaffected by IGF-2 treatment. **(B)** Stride lengths were increased in mat-/pat+ mice and were further increased by IGF-2 while **(C)** the reduced stride frequency of mat-/pat+ mice was further decreased in forelimbs by IGF-2. **(D)** IGF-2 had no effect on the elevated forelimb propulsion time of mat-/pat+ mice and led to further elevation of the increased hindlimb propulsion time. **(E)** Swing time was elevated in mat-/pat+ mice, regardless of IGF-2 treatment and **(F)** brake time was normal in mat-/pat+ mice and unchanged by IGF-2. Data are expressed as mean ± S.E.M. *n*=17-24 mice/group. A- F: \**p*<0.05, Sidak’s or Tukey’s multiple comparisons test following two-way ANOVA (per limb set). ns, not significantly different, *p*>0.05.

## Discussion

Novel data uncovered by this work illustrated that acute systemic administration of IGF-2 reduced delta spectral power in EEG, a theorized biomarker in AS. This was a very promising initial finding, considering newly published data linking delta power to improvements in the Bayley Cognitive Assessment [67], however, disappointingly, the overwhelming majority of metrics for motor behavior, learning, and coordination were unaffected and IGF-2 did not improve pup social communication, seizure threshold, cognition, or gait. Although our study returned mostly negative results regarding the potential for IGF-2 to improve behavioral deficits in AS, our findings are nevertheless important to disseminate, as they contrast other reports [46]. While we were aiming to corroborate the previous reports of IGF-2 efficacy, as inter-laboratory reproducibility is a long-standing goal of ours, we did establish strong reproducibility with other rat studies [16, 68–70], EEG and sleep studies [69, 71–74], and other genetic mutant mouse models of neurodevelopmental disorders [48, 53, 75]. Furthermore, we did reproduce a number of the *Ube3a*^mat-/pat+^ mouse phenotypes observed by Cruz et al., specifically hypolocomotion, fewer marbles buried, and poor rotarod performance [46].

We observed a moderate effect of IGF-2 on day 1 of rotarod testing in *Ube3a*^mat-/pat+^ mice, but this did not extend across the rotarod time course that addresses motor learning and it was non-existent in *Ube3a*^mat-/pat+^ rats. However, we were able to replicate all of the *Ube3a*^mat-/pat+^ mouse and rat model deficits previously reported by our groups [16, 56, 72, 76] and discover significant reduction of the elevated delta power in *Ube3a*^mat-/pat+^ EEG by IGF-2 treatment. This is the first report of detection of alterations in EEG power spectral density (PSD) without any behavioral phenotypic change. One potential explanation as to why we observed effects on EEG activity but no changes in behavioral performance is that the increase in delta power may not have substantial behavioral significance. To our knowledge, there is still little data showing that delta power is strongly tied to behavioral outcomes, despite many laboratories’ working hypothesis that PSDs are effective biomarkers [69, 74, 77–80]. However, we find this explanation unlikely in light of recent evidence from our laboratory illustrating reductions delta power with concomitant behavioral improvements [81] and a new report in humans with Angelman Syndrome [67].

Given that we were unable to reproduce, nor extend, the broad phenotypic rescue shown in earlier work, it is critical to highlight that our study employed standardized experimental protocols for behavioral testing [51, 82, 83], which differed from those used by Cruz et al. (2020) [46]. We had aimed to leverage these protocol differences to show that the effects of IGF-2 treatment were robust enough to carry across laboratories and therefore bode well for translation to the clinic. Inter-laboratory methodological discrepancies included rotarod inter-trial interval duration, open field lighting and duration, marble burying experimental design and analysis, as well as object exploration times and post-training delays. When observing latencies, we did not record scores that exceeded the duration of the test (e.g., Figure 4, Cruz et al., 2020). Additionally, while our washout period was shorter compared to previous work, we do not suspect that this hindered our ability to detect effects of IGF-2 since we did not find evidence of IGF-2 having an effect greater than one day in duration. Furthermore, if our washout period had been inadequate, the compounding effects of IGF-2 would have been revealed in subsequent testing. However, this was not the case and for each cohort of animals tested, the final assay of the test battery revealed no effect of IGF-2. Arguably, one of the most crucial methodological details that sets our behavioural experiments apart from those conducted previously is our large sample sizes, which were upwards of 25 animals per group. Pooling data from small subgroups (i.e., *n*=3-4/group as used by Cruz et al.) can artificially inflate error rates (i.e., produce false positives and negatives) due to the high risk of “testing until significance,” particularly when group sizes are not pre-determined [50, 51, 54, 83]. Pooling subgroups also requires that all groups be subjected to the same exact conditions (e.g., same sequence of prior tests, identical test parameters) and that scores from the various subgroups (particularly wildtype) are confirmed to be similar to each other. Rather than subgroups, it is recommended practice in rodent behavioral testing to use full groups consisting of 10 to 20 animals for a given experiment [51, 83]. We, therefore, only used small groups in the collection of initial pilot data and we used large cohorts with enough subjects per group to achieve robust statistical power for collection of behavioral data. The novel object recognition findings in the prior report utilized a protocol which i) we used in congenic B6J mice but were unable to reproduce previous results (i.e., there was not recognition as defined by greater time spent with novel vs. familiar object) and ii) does not appear congruent with many of the best recommended practices disseminated by the IDDRC behavioral working group (e.g., maximizing experimenter consistency, ensuring no intrinsic object preference, and using new object pairs when re- testing animals) [84].

Our study was thorough and unique, as we used two different model species and statistically powerful, large sample sizes, and we investigated the strongest reported phenotypes in the established models. Our dual species approach allowed us to measure social communication in the rat, which exhibits more nuanced social behavior and employs a more sophisticated communication system as compared to the mouse, and we leveraged the mouse model for its strong motor phenotypes. Because our rotarod paradigm consisted of three consecutive days, we were able to assess motor learning and not just use it to test motor function. Having both of these metrics available in both species was key as wildtype mice exhibited a ceiling effect that impeded interpretation of a motor learning deficit, but we were able to evaluate this outcome in rats since their performance changed significantly across test days. By comparing results across species, and across tests within the same behavioral domain, we are able to provide a more thorough and convincing assessment of this IGF-2 treatment paradigm.

While we did see a few promising trends in EEG and rotarod, we also detected effects on gait in the opposite direction than desired (i.e., worsening the phenotype), and the overwhelming majority of our findings indicate that any effect of IGF-2 is minor and does not lead to robust, reliable, or reproducible behavioral changes in either genotype. Moreover, IGF-2 treatment did not lead to consistent phenotypes in the previous report by Cruz et al. (2020). For instance, IGF-2 was not found to affect motor activity in an open field but it did lead to increased marble burying, despite motor playing a key role in marble burying behavior. We did not observe alterations in wildtype mice, which suggests that IGF-2 does not have motor, communication, or cognition enhancing properties in the time windows we assessed. Furthermore, we did not observe alteration in seizure threshold or susceptibility. Obvious differences were Cruz et al.’s utilization of 129 background mice for their audiogenic seizure procedure. AS model mice on the traditional B6J background do not exhibit spontaneous seizures nor susceptibility to audiogenic seizures [66]. We utilized the B6J background with a chemo-convulsant as 129s have a 70% reduction in corpus callosum volume which adds to their seizure susceptibility [79, 80], and sensory-dependent audiogenic seizures are triggered by divergent neural circuitry compared to chemo-induction [81].

Therapeutic mimetics of the IGF pathway are being evaluated as small molecule therapy for AS. They activate PI3K-Akt-mTOR and Ras-MAPK-ERK pathways and have been shown to increase synapse number and synaptic plasticity [85, 86]. Spine numbers have been shown to be reduced in AS mouse models [87] and activity dependent ERK phosphorylation and synaptic plasticity are impaired [88–91]. The therapeutic hypothesis is that through upregulating synaptic plasticity and synapse number, these compounds may have benefit in AS. We wanted to disseminate our mostly negative data as cautionary for interpreting IGF-2 data, as this ligand shows some non-specificity in binding both the IGF-1 and IGF-2 receptors. IGF-1 is currently being pursued as a treatment for neurodevelopmental disorders via four clinical trials: pilot clinical studies of IGF-1 are being conducted in non-genetically specified autism (NCT01970345); two clinical studies of IGF-1 are in process for Phelan McDermid Syndrome, which is a rare genetic neurodevelopmental disorder associated with mutations in *SHANK3* and one of the most common comorbid autism-associated syndromes (NCT01970345; NCT04003207), accounting for up to ∼1 of all syndromic autism [92, 93]; and clinical testing of IGF-1 in Rett Syndrome is also ongoing (NCT01777542).

## Limitations

The major limitation of the present study is that the results are confined to the three doses (10, 30, and 60 µg/kg) and one route of administration (acute subcutaneous injection) used. Particularly, our behavioral results are limited to a 30 µg/kg injection of IGF-2 delivered 20 min prior to behavioral testing. It remains possible that different doses, injection timing and/or frequency, post-administration interval, and/or routes of administration may show greater efficacy in improving the endpoints measured herein. For instance, our negative results using an acute systemic treatment of IGF-2 do not preclude the possibility that chronic delivery of IGF-2 could ameliorate behavioral deficits over longer periods of time. Additionally, our investigation of learning and memory phenotypes was relatively limited so future work would be required to comprehensively determine whether IGF-2 could ameliorate learning and memory deficits.

## Conclusions

IGF-2 did not show robust effects on key behavioral domains of relevance to AS in two genetic rodent models of AS, in contrast to a recently published report. Our findings are cautionary and emphasize that it is important for separate labs to try to replicate each other’s experiments – after all, we are in pursuit of therapeutics with broad and robust efficacy that stand up to the test of minor cross-lab methodological variations. Minimally two cohorts with standardized methods from the literature should be evaluated. Future studies that examine EEG activity during behavioral tasks may be the most informative to confirm that subtle alterations in spectral power have functional meaning before its confirmation as a robust biomarker.

## List of additional files

File name: Supplementary File 1

File format: .docx

Title of data: Supplementary Information

Description of data: Supplementary methods and figures

File name: Supplementary File 2

File format: .xlsx

Title of data: Supplementary File of Statistics

Description of data: Detailed statistical parameters for each figure

## Supporting information

Supplemental Stats Table

## Abbreviations

AS: Angelman Syndrome
ASD: autism spectrum disorder
Ube3a: ubiquitin protein ligase E3A
IGF-2: insulin-like growth factor-2
USV: ultrasonic vocalization
EEG: electroencephalography
NOR: novel object recognition
PSD: power spectral density

## Declarations

### Ethics approval and consent to participate

All animal experiments were conducted in compliance with the Institutional Animal Care and Use Committee of University of California Davis or Baylor College of Medicine.

### Authors’ contributions

ELB and SPP carried out the behavioral experiments and subsequent analyses. HAB performed the electrophysiology experiments and data analysis. AA collected and analyzed behavioral data. AEA and JLS supervised the study and interpretations of data. ELB, SPP, and JLS drafted the initial manuscript. All authors made valuable comments and edits to the manuscript and approved the final version.

### Consent for publication

Not applicable.

### Availability of data and materials

The datasets used in the current study are available from the corresponding author upon request.

## Acknowledgements

We thank Annuska Berz, Markus Wöhr, Timothy Fenton, and Yutian Shen for their support with this project.

