## Supplemental Information and Figures for "Insulin-like growth factor-2 does not improve behavioral deficits in mouse and rat models of Angelman Syndrome"

**Methods**

***Novel object recognition (NOR) in C57BL/6J mice.*** Learning and memory were tested by individually presenting subjects with two identical objects and later testing their ability to recognize the familiar object over a novel one following a protocol previously described by Cruz et al. (2020) [1]. The NOR assay was carried out within an opaque matte white arena (41 cm l x 41 cm w x 30 cm h) in a 30-lux room and consisted of four phases: a 5-min habituation to the arena on the day prior to the test, a 3-min object familiarization session, a 24-hr isolation period, and a 5-min object recognition test. Twenty min prior to the familiarization phase, mice were administered 30 µg/kg IGF-2 or vehicle via subcutaneous injection. Following the 5-min habituation period, each animal was removed from the arena and placed in an individual clean holding cage while two clean identical objects were placed inside the arena. Each subject was then returned to its arena and allowed to freely explore and familiarize with the objects for 3 min. After 24 hrs, subjects were returned to their arenas and allowed to freely explore one familiar and one novel object for 5 min. Time spent investigating each object was measured manually by a trained observer blinded to treatment group and genotype. Recognition memory was defined as spending significantly more time investigating the novel object compared to the familiar object by paired *t*-test within group. Object preference was calculated as time spent sniffing the novel object compared to total time sniffing both objects. Fifty percent represents equal time investigating the novel and familiar object (a lack of preference) whereas >50% demonstrates intact recognition memory. The effect of IGF-2 on percent preference was analyzed using unpaired *t*-test (**Supplementary File 2**).

***Contextual fear conditioning in C57BL/6J mice.*** Contextual fear conditioning was carried out using an automated fear conditioning chamber (Med Associates, Inc., Fairfax, VT, USA) following the training protocol previously described by Cruz et al. (2020) [1]. On the training day, mice were administered 30 µg/kg IGF-2 or vehicle via subcutaneous injection and, after a 20 min delay, exposed to an unsignaled foot-shock within a testing chamber with specific visual, odor, and tactile cues. After 2 min, a 2-sec foot shock (0.7 mA) was delivered. A 1 min exploration period followed the noise-shock pairing before the mouse was placed back in the home cage. Twenty four hours later, the subject was placed back inside the training environment for 5 min. The chamber contained identical contextual cues as the training session, but no foot shock occurred, and the percent time spent freezing was automatically measured by VideoFreeze software (Med Associates). The effect of IGF-2 on percent time freezing was analyzed using unpaired *t*-test (**Supplementary File 2**).

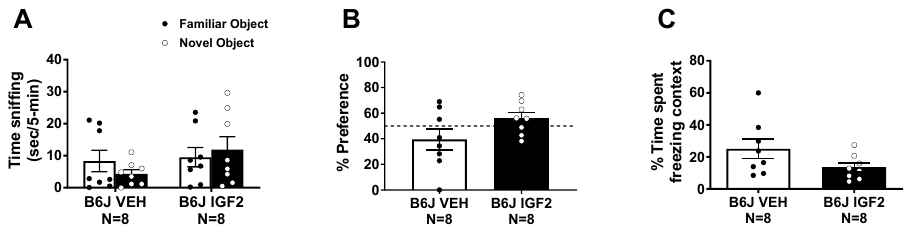

**Fig S1. IGF-2 did not enhance cognition in novel object recognition or contextual fear conditioning tasks in pure congenic C57BL/6J mice.** The cognitive enhancing capabilities of IGF-2 were assessed in congenic C57BL/6J **(**B6J) mice. **(A)** Novel object recognition was tested using the protocol of Cruz et al. (2020) with administration of 30 µg/kg IGF-2 20 minutes before the familiarization phase and the recognition memory test 24 hours after familiarization. Neither vehicle (VEH) nor IGF-2-treated mice met the criteria for recognition memory as they did not spend more time sniffing the novel object compared to the familiar object. **(B)** There was no difference between vehicle and IGF-2 groups in percent preference for the novel object. **(C)** Contextual fear conditioning was evaluated using the training protocol of Cruz et al. (2020) with administration of 30 µg/kg IGF-2 20 minutes before the training session and the contextual memory test 24 hours after training. There was no difference in percent time freezing between the vehicle and IGF-2 group. Data are expressed as mean ± S.E.M. *n*=8 mice/group.
